# Prime-boost vaccinations with two serologically distinct chimpanzee adenovirus vectors expressing SARS-CoV-2 spike or nucleocapsid tested in a hamster COVID-19 model

**DOI:** 10.1101/2022.03.17.484786

**Authors:** Mohadeseh Hasanpourghadi, Mikhail Novikov, Robert Ambrose, Arezki Chekaoui, Dakota Newman, Jianyi Ding, Wynetta Giles-Davis, Zhiquan Xiang, Xiang Yang Zhou, Qin Liu, Kar Swagata, Hildegund CJ Ertl

## Abstract

Two serologically distinct replication-defective chimpanzee-origin adenovirus (Ad) vectors (AdC) called AdC6 and AdC7 expressing the spike (S) or nucleocapsid (N) proteins of an early SARS-CoV-2 isolate were tested individually or as a mixture in a hamster COVID-19 challenge model. The N protein, which was expressed as a fusion protein within herpes simplex virus glycoprotein D (gD) stimulated antibodies and CD8^+^ T cells. The S protein expressing AdC (AdC-S) vectors induced antibodies including those with neutralizing activity that in part cross-reacted with viral variants. Hamsters vaccinated with the AdC-S vectors were protected against serious disease and showed accelerated recovery upon SARS-CoV-2 challenge. Protection was enhanced if AdC-S vectors were given together with the AdC vaccines that expressed the gDN fusion protein (AdC-gDN). In contrast hamsters that just received the AdC-gDN vaccines showed only marginal lessening of symptoms compared to control animals. These results indicate that immune response to the N protein that is less variable that the S protein may potentiate and prolong protection achieved by the currently used genetic COVID-19 vaccines.

## INTRODUCTION

SARS-CoV-2 was first detected in humans towards the end of 2019. Since then, it has infected over 400 million humans and killed nearly 6 million. Although vaccines that are highly efficacious against disease caused by the initial isolates have been available for over a year, the virus is constantly evolving and one of the latest variants, called omicron, in part evades vaccine-induced virus neutralizing antibodies (VNAs) [1].

Vaccines that are currently available are either genetic vaccines in form of mRNA vaccines[2,3], Ad vector vaccines [4–6], or they are based on protein vaccines [7] or inactivated viral vaccines [8]. Vaccines but for those using inactivated virus express only the SARS-CoV-2 S protein, which is the target antigen for VNAs [9] that were shown by passive transfer experiments to protect against an infection [10]. Two doses of mRNA vaccines were found to be initially over 90% effective in preventing symptomatic disease [2,3]. The Ad vector-based Sputnik V vaccine given twice in a heterologous prime boost regimen is equally protective^6^ while the single dose Ad vector vaccine from Johnson and Johnson (J&J) [4] and the AdC vector vaccine from AstraZeneca that uses the same vector for priming and boosting protect 63-67% of individuals against disease [5], which is similar to the level of protection achieved with the protein vaccine^7^ or the inactivated viral vaccines [8]. As has been studied most extensively with mRNA vaccines efficacy is not sustained necessitating booster immunizations after 6 months [11,12]. Notwithstanding, even fully vaccinated individuals experience breakthrough infections [13,14] especially with omicron [15] and although their symptoms are in general mild or nonexistent, they can spread the virus to others. SARS-CoV-2 will continue to evolve and infections of vaccinated or previously infected individuals will favor outgrowth of mutants that may escape vaccine-induced VNAs even further.

T cells induced by natural infections with common cold coronaviruses can cross-react with SARS-CoV-2 antigens [16] and they have been implicated to provide protection to so-called ‘never COVID’ individuals, who despite close contacts with SARS-CoV-2-infected individuals remain uninfected [17]. The N protein is one of the most abundantly expressed viral antigen in SARS-CoV-2 infected cells, which should make it an easy target for specific T cells. Antibodies to the N protein may also contribute to viral clearance through antibody dependent cellular cytotoxicity (ADCC) or complement-mediated lysis. N proteins potential role as a vaccine antigen to induce sustained and cross-reactive immunity to SARS-CoV-2 warrants further investigations.

This was the goal of our study, which tested in a hamster challenge model, vaccines that not only express the S protein for induction of VNAs but also the more conserved N protein. We expressed each of the two SARS-CoV-2 proteins by two serologically distinct replication-defective AdC vectors of serotype SAdV-26, also called AdC6, and SAdV-24, also called AdC7 [18,19]. Inserts were based on unmodified genes from an early viral isolate from Sweden, called SARS-CoV-2/human/SWE/01/2020 (AdC-S_SWE_, GenBank number: QIC53204), whose S and N protein sequences are identical to those of the original Wuhan isolate. The N protein was expressed as a fusion protein within herpes simplex virus gD, which through inhibition of the early B and T cell attenuator (BTLA) -herpes virus entry mediator (HVEM) checkpoint broadens SARS-CoV-2 N protein-specific CD8^+^ T cell responses [20,21] and augments B cell responses [22]. Vaccines were given in a heterologous prime boost regimen either individually or as mixtures of the same vector backbones expressing the two different inserts. Our data show that vaccines that express the gD-N fusion protein induce N-protein-specific antibody and CD8^+^ T cell responses but provide robust protection against infection or disease. As expected, the S protein-expressing vaccines induce VNAs and reduce disease after SARS-CoV-2 challenge but fail to prevent infection. Best protection was achieved with the vaccine mixtures indicating that although N protein-specific immune mechanisms by themselves fail to lessen disease upon challenge with a high dose of SARS-CoV-2 they enhance protection mediated by S protein-immunity.

## RESULTS

### Experimental design

Sixteen female and sixteen male Syrian golden hamsters were enrolled and separated into 4 groups of 8 animals each (4 females, 4 males/group). The groups were primed by intramuscular (i.m.) injections as follows: group 1 was injected with 1 × 10^10^ virus particle (vp) of AdC7-gDN, group 2 with 1 ×10^10^ vp of AdC7-S, group 3 with a mixture of 0.5 × 10^10^ vp AdC7-gDN and 0.5 × 10^10^ AdC6-S and group 4 with 1 × 10^10^ vp of the control vector, called AdC7-HBV2, which expresses a sequence derived from hepatitis B virus. We only used half of the vaccine dose for each component of the mixture to keep the total amount of vector given to each animal constant, as in a clinical setting higher vaccine doses translate to increases in adverse events driven by innate responses to the Ad vector [23]. Animals were boosted 2 months after priming with AdC6 vectors expressing the same inserts and used at the same doses. Hamsters were bled at baseline, 14 days after the prime and 14 days after the boost to determine SARS-CoV-2 S protein-specific antibody responses in groups 2-4. N-specific antibody and CD8^+^ T cell responses in groups 1, 3, and 4 were measured from blood 14 days after the boost (Figure 1A). Animals were challenged intranasally with SARS-CoV-2 4 weeks after the boost. After challenge animals were checked daily for clinical symptoms and weight loss. Oral swaps were collected on days 2, 4, 7, and 14 after viral challenge to determine levels of viral genomic and sub-genomic (sg)RNA. Four animals in each group were euthanized 4 days and the others 14 days after challenge. After euthanasia, lung viral titers were determined, and lung sections were screened for pathology.

**Figure 1.**
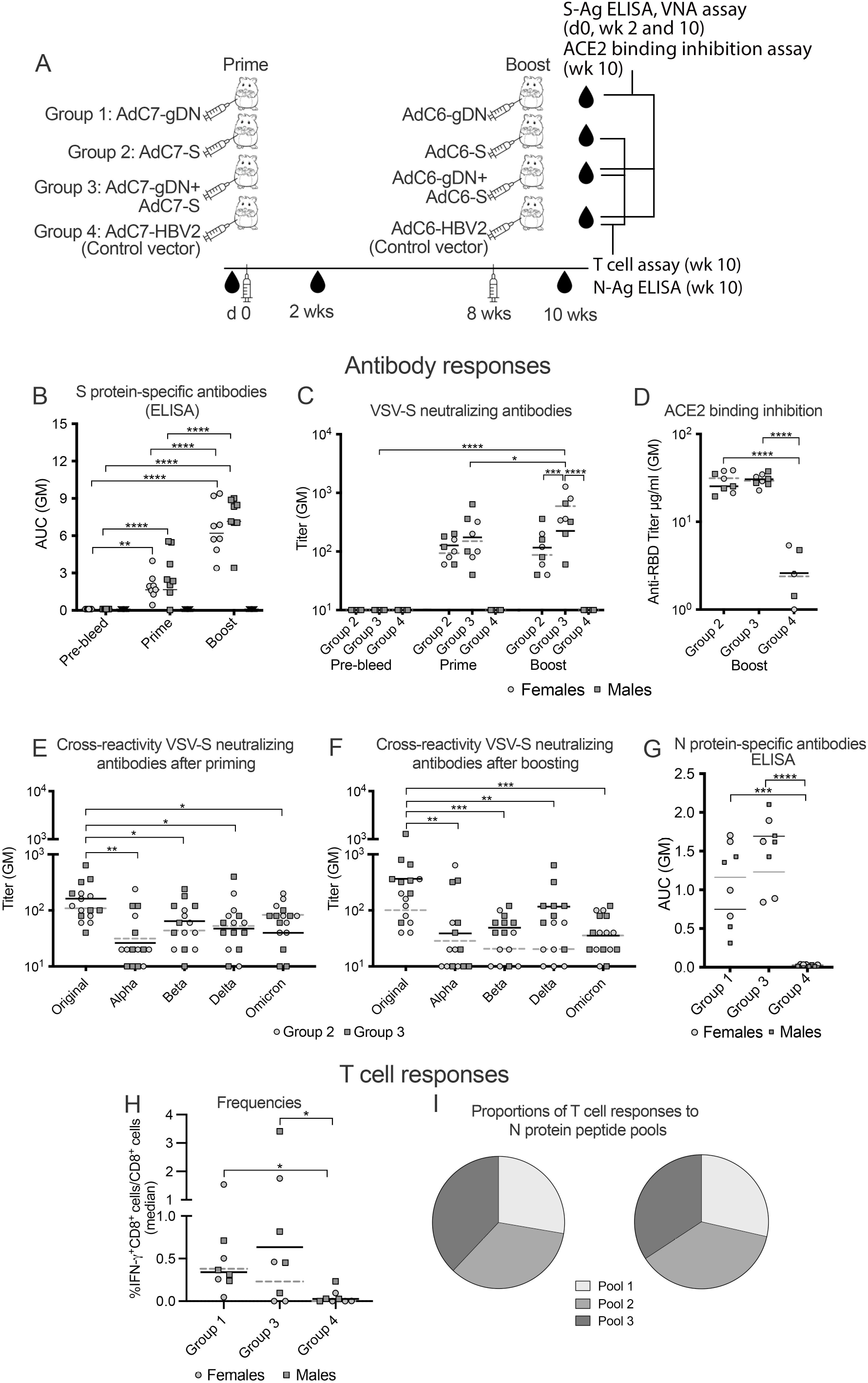
Immune responses to the vaccine vectors. [A] Experimental design. [B,C] Sera harvested at baseline, 2 weeks after the prime or the boost were tested for antibody responses. [B] Reactivity against SARS-CoV-2 S1/S2 proteins tested for by an ELISA. Data show area under the curves (AUC) for dilution curves generated with sera of individual mice. Control animals scored negative with adsorbance values below background and these data are shown as an AUC of 0.1. Lines show geometric means (GM). Significant differences were calculated by 2-way Anova with Tukey correction. In this and all subsequent graphs lines with stars above indicated significant differences: (*) p-value between 0.01-0.05, (**) p-value between 0.001 – 0.01., (***) p-value between 0.0001-0.001, (****) p-value <0.0001. [C] Sera were tested by a neutralization assay using VSV-S vectors pseudotyped with the same S protein as in the vaccines. Data are shown for individual animals. Negatives are shown as a titer of 10. Lines indicate GMs. Lines with stars above show significant differences by 2-way Anova. [D] Sera harvested after the boost were tested for inhibition of ACE binding to the RBD of S1 by an ELISA. Data are shown for individual mice as µg of antibody/ml calculated based on an internal standard. Negatives are shown as a titers of 1 µg/ml. Lines indicate GMs. Significant differences were calculated by Kruskal Wallis test. [E] [E,F] Cross-reactivity against SARS-Co-V2 variants. Sera collected after the prime [E] or the boost [F] were tested for neutralization of VSV-S vectors pseudotyped with the indicated S protein variants. Group 2 [light grey circles] and group 3 [dark grey squares] are indicated by different symbols. For the statistical analysis by one-way Anova data for the two groups were combined. [G] Sera harvested 2 weeks after the boost were tested for N protein-specific antibodies by an ELISA. Results are shown as AUC for individual sera from female and male hamsters. Lines show GMs. Significant differences were calculated by 2-way Anova with Tukey correction. [H] Frequencies of CD8^+^ T cells in blood producing IFN-γ in response to N-derived peptides. Graph shows sum of responses to the 3 peptide pools. Lines show median responses. Lines with stars above show significant differences between animals that received the COVID vaccine and control animals by Kruskal Wallis test. [I] Proportions of N-specific CD8^+^ T cell responses to the 3 peptide pools. [B-E] Results for females (light circles) and males (darker squares) are indicated by the different symbols.

### Immune responses to the vaccine vectors

Sera from hamsters of groups 2, 3, and 4 harvested at baseline and at 2 weeks after the prime or boost were tested by ELISA on plates coated with S1 and S2 proteins, which measures all S-binding antibodies. They were also tested in a neutralization assays using vesicular stomatitis virus (VSV) vectors pseudo-typed with SARS-CoV-2 S protein to detect VNAs, which can either be directed to the RBD sequence of S1 or the fusion peptide in S2. Sera harvested 2 weeks after the boost were in addition tested by an ELISA, which measures antibodies specific to the receptor binding domain (RBD) of S1 (Figure 1A). All pre-immunization samples and samples from the control groups harvested at either time point scored negative in either one of these assays. All but one of the sera from one hamster of groups 2 scored positive by the S1/S2-specific ELISA by 2 weeks after the prime. All hamsters seroconverted after the boost and overall responses showed significant increases (Figure 1B). More specifically geometric mean (GM) titers of group 2 females increased 1.6-fold, while those of males increased 6.8-fold; GM titers of group 3 females increased 1.5-fold while those of males increased 1.9-fold. There were no significant differences in responses of males compared to females although in group 2 males tended to have higher responses after the boost than females. All hamsters developed antibodies that neutralized the pseudotyped VSV-S virus with GM titers of 109 in group 2 and 167 in group 3 after the prime which changed to 166 and 365 after the boost, respectively. The superior booster response in group 3 was driven by female hamsters, which showed a 4-fold increase in VNA titers compared to the marginal 1.2-fold increase in males. After the boost all animals scored positive in the RBD-specific ELISA (Figure 1D) and there were no significant differences between males and females or groups 2 and 3. VNAs were tested for reactivity against alpha, beta, delta, and omicron variants. Sera collected after the prime (Figure 1E) or after the boost (Figure 1F) showed significantly reduced titers comparing neutralization of VSV-S vectors expressing wild-type S protein to those with S proteins of variants.

Sera from hamsters of groups 1, 3, and 4 harvested after the boost were tested for antibodies to the N protein. All AdC-gDN vaccinated animals scored positive with no differences between groups 1 and 3 or males and females hamsters (Figure 1G).

Peripheral blood lymphocytes were tested two weeks after the boost for production of IFN-γ by CD8^+^ T cells in response to three peptide pools representing the N protein sequence (Supplemental Table 1). Most animals of groups 1 and 3 developed N protein-specific CD8^+^ T cell responses; 1 and 2 females of groups 1 and 3, respectively and 1 male of group 3 failed to respond (Figure 1F). Responses showed no preference to any of the 3 peptide pools representing different segments of the N protein.

### Clinical symptoms and weigh loss after SARS-CoV-2 challenge

Four weeks after the boost animals were challenged intranasally with SARS-CoV-2 (Figure 2A). Animals were scored twice daily for clinical symptoms. Most animals developed benign symptoms after challenge in form of mildly ruffled fur, hunched over posture and/or closed or squinted eyes. None of the vaccinated mice exceeded a COVID-19 disease score of 1 and only one female in the control group 4 was given a disease score of 2 on days 5-7 after challenge. One male animal in the control group died 7 days after challenge although it never exceeded a disease score of 1. His lung pathology suggested that his death was most likely caused by SARS-CoV-2.

**Figure 2.**
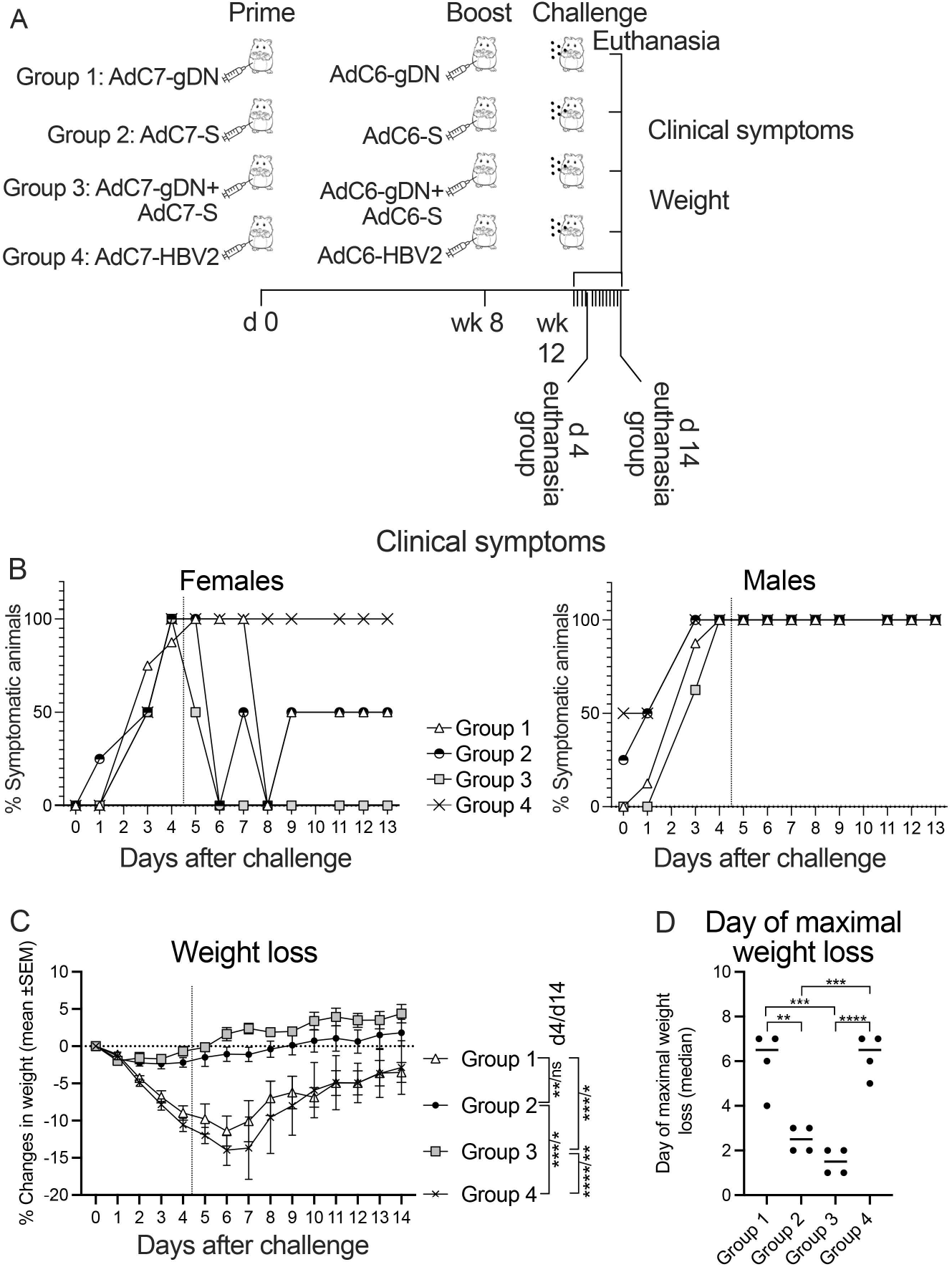
Disease score and weight loss after challenge. [A] Experimental design. [B] % of female (left) and male (right) animals that scored positive for disease on the indicated days after challenge. Morning and afternoon scores were averaged. [C] Weight loss after challenge is shown as % weigh reduction over the weight on the day of challenge. Data are shown as means ± SEM. Significant differences calculated by two-way Anova with Tukey correction for day 4 and 14 data are indicated by connecting lines next to the legend. Day 4 comparisons are shown first followed by / and then day 14 comparisons.

Plotting days of clinical symptoms against days after challenge showed that most female animals developed symptoms by day 4 after challenge but for one group 3 animal, which became symptomatic on day 5. All males of group 4 and most of the males of the vaccine groups had symptoms by day 3 and by day 4 all male animals had visible signs of disease. Thereafter the disease course was markedly different for vaccinated female and male animals as has also been reported for humans [24]. In both gender groups unvaccinated hamsters exhibited symptoms till the day of their euthanasia. The same was seen for all males of the vaccine groups. With some fluctuations half of the females of group 1 and 2 remained symptomatic throughout the observation period while group 3 females became and remained symptom-free by days 5 or 6 (Figure 2B). Although this result is based on small numbers of animals is nevertheless suggests that vaccinated females recover more rapidly than vaccinated males upon breakthrough infections and that the combination vaccine compared to the individual vaccines accelerated recovery in females.

These data were mirrored by levels of weight loss. Control animals rapidly started losing weight after challenge (Figure 2C) with maximal weight loss around days 6-7 (Figure 2D). Group 1 animals which had only received AdC-gDN vectors lost weight with similar kinetics to the control group although their maximal weight loss by days 6 and 7 after infection tended to be lower. Hamsters of groups 2 and 3 lost only a minimal amount of weight early after challenge and started gaining weight by day 2 or 3 after challenge. On days 4 and 14 after challenge groups 2 and 3 showed significantly less weight loss compared to the control group 4; group 3 showed a significant difference to group 1 at both time points while group 2 only differed from group 1 by day 4 after infection. Females recovered their weight 1-2 days earlier than males (Suppl. Figure 1A).

### Effect of SARS-CoV-2 on viral loads and lung histology

Viral titers were determined from oral swabs collected on days 2, 4 (8 animals for each group), 7 and 14 (4 animals for each group) (Figure 3A) and lung tissues collected from 4 animals of each group either on days 4 or 14 after challenge. Samples were screened for viral RNA and sgRNA, the latter has been suggested to determine titers of infectious virus more accurately [25]. Immunization with the vaccine mixture reduced oral viral RNA (Figure 3A) and sgRNA (Figure 3B) titers on day 2 after infection while immunization with the Ad-S vaccine only reduced sgRNA titers but not RNA titers on day 2. No significant differences were observed for the group 1 or the other time points. Vaccination with either the AdC-S or the AdC-S + AdC-gDN regimens reduced lung viral RNA titers (Figure 3C) tested on day 4 after challenge while AdC-S vaccination also reduced sgRNA titers at that time point (Figure 3D). By day 14 viral loads in lungs were markedly reduced in all groups and although group 3 continued to show lower RNA and sgRNA titers compared to the others this failed to reach significance.

**Figure 3.**
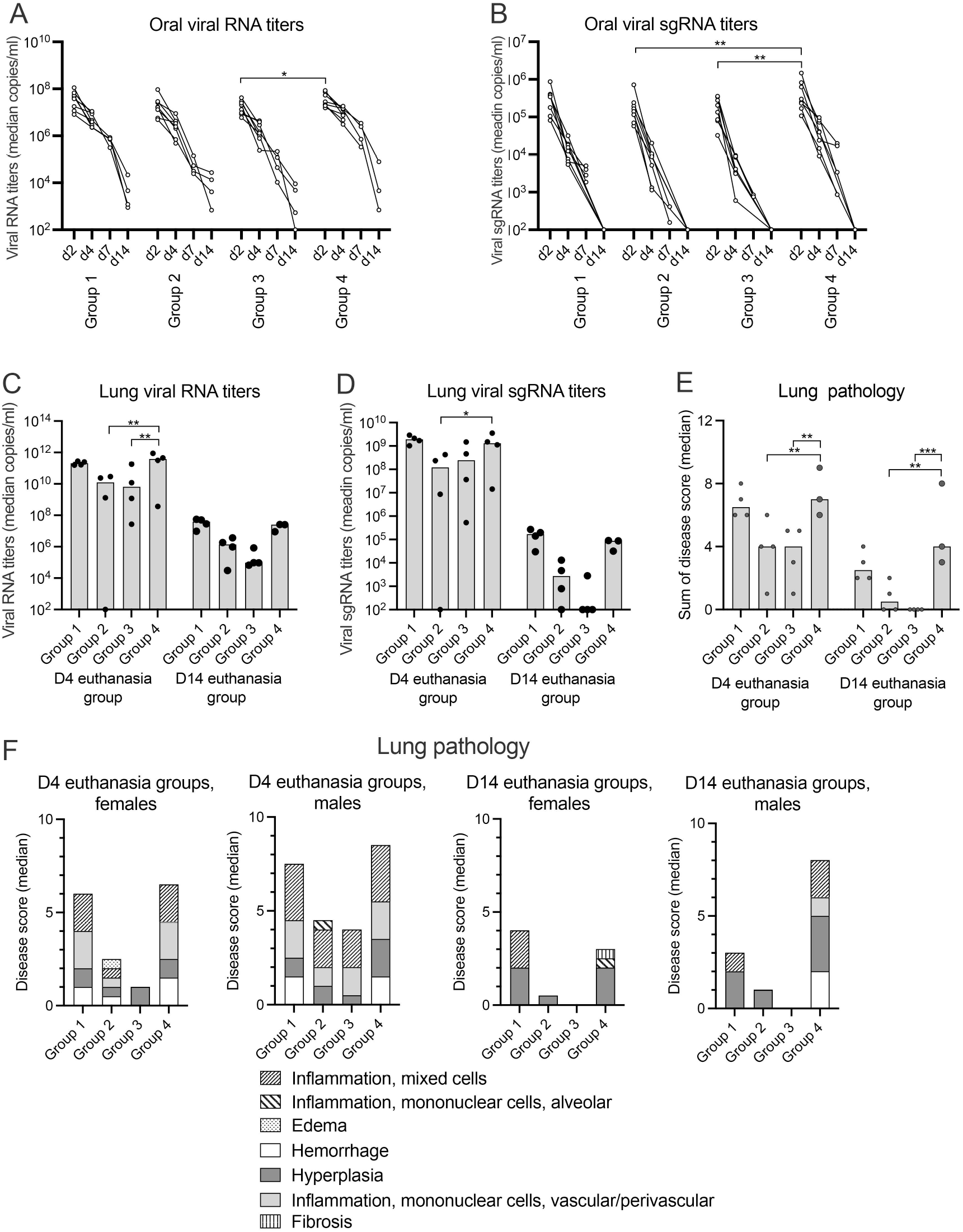
Viral loads and lung pathology. [A,B] Viral RNA [A] and sgRNA [B] loads in oral swabs collected after challenge on days 2 and 4 for all animals and on days 7 and 14 for the day 14 euthanasia group animals. Significant difference for each time point were calculated by 2-way Anova ; they are indicated with lines and stars above as in legend to Figure 1. [C,D] Lung viral RNA [C] and sgRNA [D] titers are shown for individual hamsters of the day 4 and day 14 euthanasia groups. Differences were calculated by 2-way Anova with lines and stars showing significant differences as in legend to Figure 1. [E] Sum of lung lesions for the day 4 and day 14 euthanasia group. Differences were calculated by 2-way Anova with lines and stars above as in legend to Figure 1. [F] Severity of the different types of lesions according to gender.

Parts of the lungs were sectioned and stained after euthanasia for 4 animals of each group on day 4 and the remaining animals on day 14 after challenge and analyzed for pathological changes (Figure 3E). Macroscopically lungs showed red or dark red patches, dark spotted, mottled/red lobes in groups 1, 2, and 4 but not 3 (Suppl. Figure 1B). Tissue lesions were graded microscopically and lungs from most animals showed grade 1-3 histopathology. Especially on day 4 lungs showed perivascular edema, alveolar hemorrhage, bronchiolo-alveolar hyperplasia, mixed or mononuclear cell inflammation (bronchoalveolar, alveolar, and/or interstitial), mononuclear vascular/ perivascular inflammation, mesothelial hypertrophy, and/or pleural fibrosis (Figure 3E,F). In groups 2 and 3, lesions were milder and more multifocal than in groups 1 and 4 on day 4. The sum of the score for the individual symptoms showed on day 4 significant differences between groups 2 and 3 compared to the control animals of group 4. On day 14 lungs of the control animals continued to show marked pathology while group 3 animal lungs no longer showed any lesions; group 2 animals showed residual mild disease. Group 1 animals, which only recieved the AdC-gDN vaccine, also showed reduced pathology compared to group 4 animals (Figure 3E). Lung pathology showed gender specific differences (Figure 3F). By day 4 after infection the two group 3 females only showed grade 1 hyperplasia while the 2 males of this group had additional lesions. By day 14 both females of the control group showed a lessening of symptoms, which was not observed in the surviving male.

### Correlations

We analyzed the data for correlations by Spearman separately for hamsters that were tested for S protein specific antibody responses (groups 2 and 3) and/or N protein specific T and B cell responses (group 1, 3, and 4). T cell responses inversely correlated with oral RNA and sgRNA titers on day 2 (p = 0.037/0.02) while S protein specific antibody by the S1/S2 ELISA after the boost showed inverse correlations for oral viral RNA or sgRNA titers on day 4 (ELISA: p = 0.049/0.029). For other parameters such as relative weight after challenge, lung pathology or lung viral titers we analyzed animals that were euthanized on day 4 separately from those that were euthanized on day 14 again keeping the T and B cell groups separate. These analyses showed as expected direct correlations between lung viral loads and lung pathology. We did not observe significant correlations between T cell responses and any of the parameters of disease. In the early euthanasia group VNA titers after the boost inversely correlated with RNA and sgRNA lung viral titers on day 4 (RNA p = 0.01, sgRNA p = 0.006) while the late euthanasia group showed antibody responses by S1/S2 ELISA after the boost inversely correlated with oral viral RNA or sgRNA titers on day 4 (p = 0.01 for both). The most pronounced correlations were seen for antibody responses to the N proteins in groups 1, 3, and 4. N protein specific antibody responses showed for the analyses with all animals and the day 4 or 14 euthanasia group animals strong positive correlations with relative weight and for the former two analyses strong inverse correlations with RNA and sgRNA titers in saliva on day 4. The day 14 euthanasia animals also showed strong inverse correlations between N protein specific antibody titers and sgRNA and RNA titers in saliva on days 2 and 4, lung sgRNA titers and lung pathology (Figure 4).

**Figure 4.**
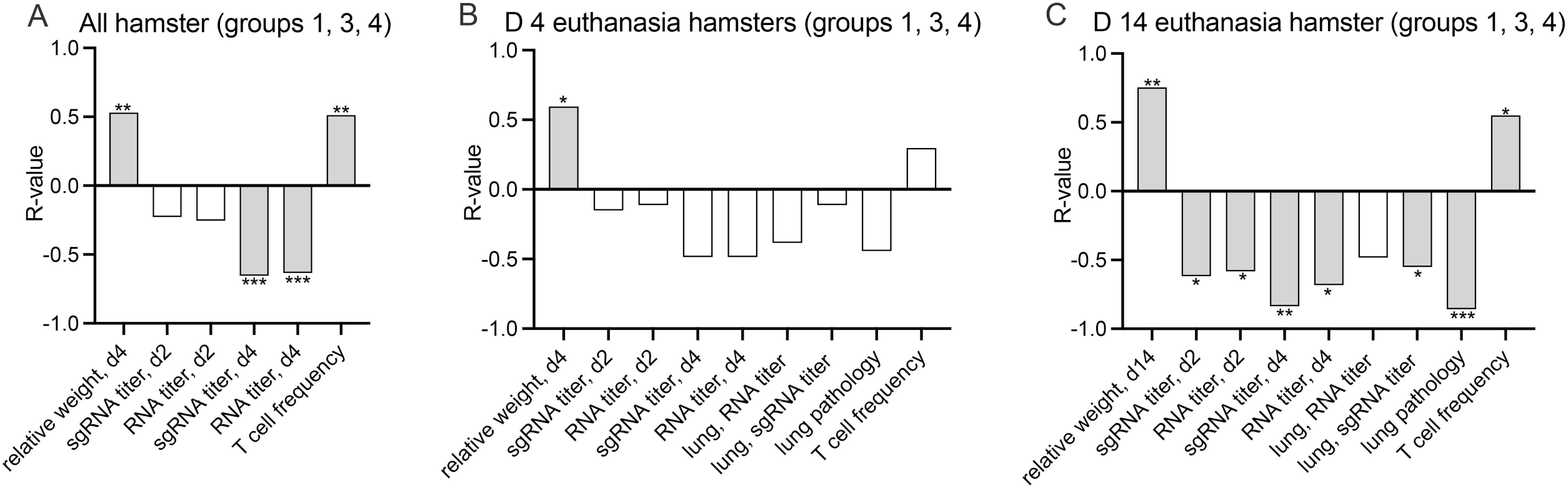
Correlations between antibodies to N proteins and disease parameters and T cell responses. The graphs show the R-values of correlations by Spearman between N protein specific antibody responses and the indicated parameters of hamsters from groups 1, 3 and 4. [A] shows the analysis for all animals. [B] shows the analysis for the day 4 euthanasia animals. [C] shows the analysis for the day 14 euthanasia animals. [A-C] Significant correlations are shown by light grey bars, data that did not reach significance are shown as white bars. Level of significance is indicated by strs above the bars as described in legend to Figure 1.

### Conservation of T and B cell epitopes

Our results show that vaccine induced S protein-specific immune responses reduce COVID-19 disease after SARS-CoV-2 infection and that addition of vaccines expressing the N protein has benefits. The S protein of SARS-CoV-2 has undergone extensive mutations, some of which increase viral transmission [26,27] and affect neutralization by vaccine-induced antibodies [28]. Both the S and N protein have numerous MHC class I and II epitopes for recognition by CD8^+^ and CD4^+^ T cells (Suppl.Figs. 2-4) and such sequences may be more conserved than those of VNA binding epitopes. Using epitope prediction software (http://tools.iedb.org/main/tcell/), we analyzed HLA class I and II epitopes present within the SARS-CoV-2 S and N protein sequences and how these epitopes have been affected by mutations within variants. For HLA class I epitopes we used a cut-off score of 0.8 while for HLA class II epitopes a cut-off rank of ≤ 1 was used. The S protein is very rich in HLA class I epitopes and about 44% of the sequence can be recognized by CD8^+^ T cells from individuals with different HLAs while 24% of the sequence are covered by HLA class II epitopes. T cell epitopes are less abundant in the shorter N protein; 36% and 11% of the sequence scored as potential HLA class I or II epitopes respectively (Figure 5A). Of the mutations within the alpha or beta variants only a modest percentage affected HLA class I epitopes while within delta and omicron mutations were present in 38% or 48% of such epitopes. HLA class II epitopes within the S protein were less affected and showed mutations rates between 13-27% for the different variants. (Figure 5B). Most of the epitope mutations within the variants resulted in a reduction or even loss of predicted binding to their corresponding restriction elements. Only one of the mutations within N present in the alpha variant affected an HLA class I epitope; this mutation resulted in increased HLA class I binding to the corresponding peptide (Figure 5C). HLA class II epitopes of N were not modified by any of the mutations (Figure 5D).

**Figure 5.**
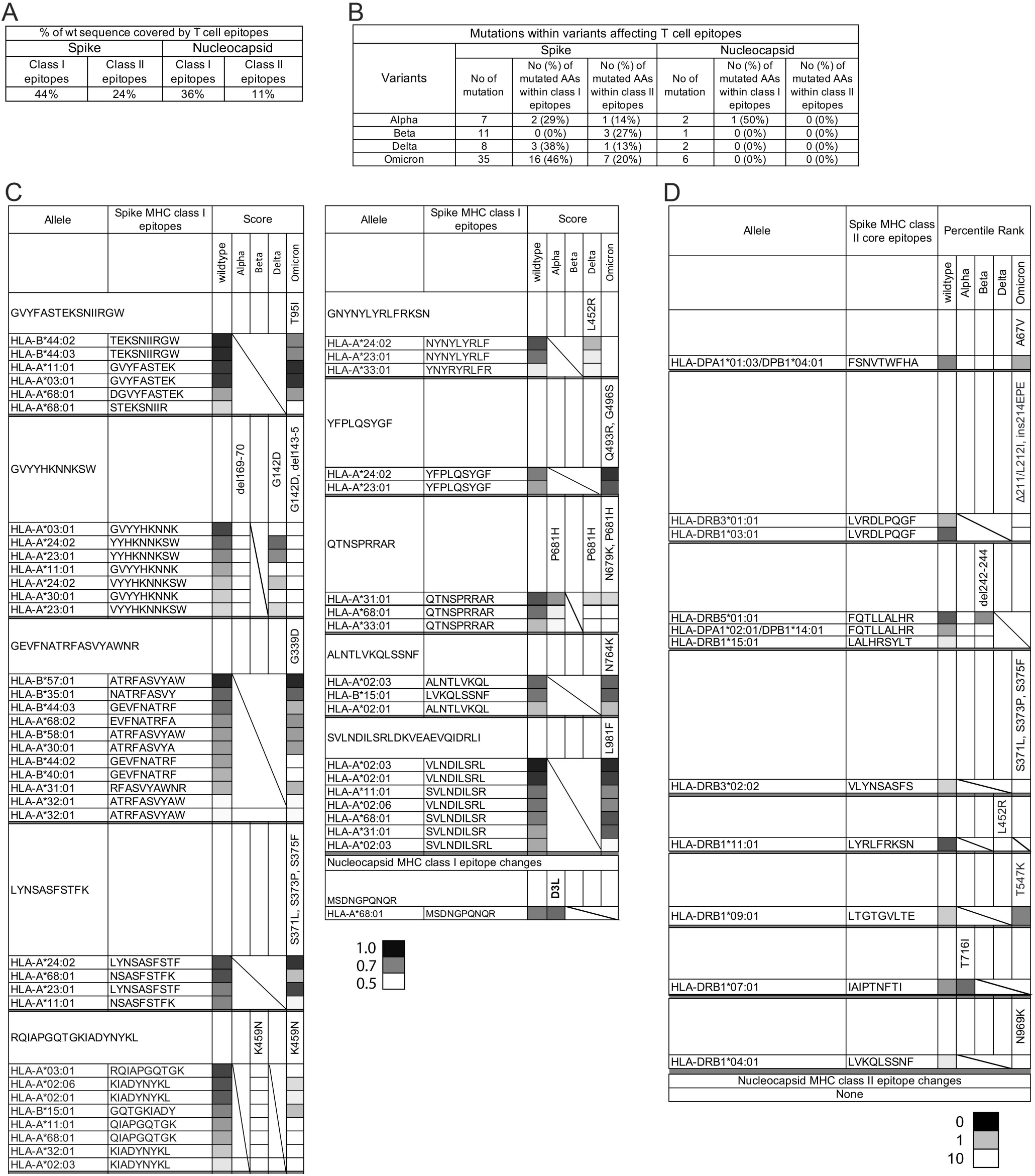
T cell epitopes within SARS-CoV-2 variants. [A] Percentage of the S and N protein sequence that is covered by putative HLA class I or II T cell epitopes. [B] Numbers of S and N protein amino acid mutations within the variants and how many of them are within T cell epitopes. [C] Different putative HLA class I epitopes and their restricting elements that are affected by mutations in the variants. The first line of each set shows to the left the AA sequence that was analyzed and to the right the mutations within this sequence. The parts below show the AA sequence that putatively binds to the HLA alleles and the left the score for the peptide sequence from the original virus (wt – wildtype) and the variants as a heatmap. [D] shows the same for HLA class II epitopes.

Several linear epitopes have been identified for SARS-CoV-2 S and N proteins [29]. Of the 4 within the S protein 3 are mutated in the variants; the 1^st^ epitope that stretches from amino acids 200-220 is mutated in all of the analyzed variants. A 2nd epitope from amino acids 544 to 562 is mutated in the alpha variant and the 3^rd^ epitope from amino acids 569 to 603 is mutated within omicron. The single linear epitope within the N protein is conserved between the variants.

## DISCUSSION

In a remarkable achievement, SARS-CoV-2 vaccines were developed and approved for use in humans within less than a year after onset of the COVID-19 pandemic [2–8]. Nevertheless, even in countries with easy access to the vaccines the pandemic has been a roller coaster ride with cycles of reductions in cases followed by rapid increases due to the emergence and spread of more transmissible variants. This constant up and down in caseloads shows how fragile our control over SARS-CoV-2 remains. Available COVID-19 vaccines protect well against disease but largely fail to prevent infections, which allows for spread of the virus even in populations with high vaccine coverage [30]. In addition, protection induced by mRNA vaccines, one of the most widely distributed types of COVID-19 vaccines, wanes after a few months. Resistance to COVID-19 disease can be restored at least temporarily by a booster immunization but after a third dose of an RNA vaccine [31], protection declines after four months [32], and it is currently unknown if frequent boosts with such genetic vaccines within short intervals will continue to restore antibody titers. Long-lived plasma cells can maintain antibody responses often for years after vaccination or infection. Induction of long-lived plasma cells required T cell help within germinal centers and their maintenance requires continuous survival signals in specialized niches of the bone marrow or secondary lymphatic tissues [33]. Although COVID-19 mRNA vaccines were shown to result in germinal center formation[34,35], available evidence including the phenotypes of the vaccine-induced B cells [35] and the rapid declines in vaccine-induced antibody titers suggest that COVID-19 mRNA vaccines induce primarily short-rather than long-lived plasma cells [36]. This combined with the unceasing emergence of new variants that more and more escape vaccine-induced VNAs necessitates the development of second generation COVID-19 vaccines that induce sustained protection against a broad spectrum of current and future variants.

SARS-CoV-2 has thus far mainly accumulated mutations within the S protein. As other viral proteins are less variable, we decided to explore vaccines that expressed the N protein in addition to the S protein vaccine. The N protein is a target for non-neutralizing antibodies that might play a role in combating a SARS-CoV-2 infection [37]. The N protein also induces CD8^+^ T cells, that by rapidly killing cells early after their infection, have also been implicated in contributing to protection against severe COVID-19 or to improve resistance in individuals with suboptimal VNA titers [38–40].

For the hamster challenge study, we used two AdC vectors that belong to distinct serotypes to prevent that Ad capsid-specific antibody responses induced by the prime neutralized the Ad vector used for the boost. Humans only rarely have VNAs to AdC viruses and those who are seropositive have low titers [41]. VNAs to Ad viruses, directed mainly against the hypervariable loops of the hexon protein [42], can prevent infection of cells with Ad vector vaccines and thereby expression of the transgene product. This in turn reduces the antigenic load and blunts immune responses to the vaccine antigen, which can be problematic for human serotype Ad vectors or for the repeated use of the same Ad vector [43,44]. The AdC vectors for the S protein carry the full-length unmodified gene from an early SARS-CoV-2 isolate and as we showed earlier induce after a boost sustained antibody responses in mice [44]. These data were recapitulated in hamsters which developed binding as well as neutralizing antibodies after vaccination with the AdC-S vectors. A booster effect was seen for binding antibodies for groups 2 and 3 while VNA responses only increased in group 3 mice that received the AdC-S/AdC-gDN vector mixtures. Whether this reflects increased T help due to responses to the N protein or a difference in vector dosing with group 2 hamsters receiving double the dose of the AdC-S vector is unknown. As expected, and reported by others, VNAs cross-reacted albeit with significantly reduced activity against viral variants of concern [45–47].

The N protein is expressed by the AdC vectors as a fusion protein within HSV-gD, which as an inhibitor of the early BTLA-HVEM T cell checkpoint enhances B cell responses and broadens CD8^+^ T cell responses as we showed with several antigens including the SARS-CoV-2 N protein in mice [20,21,38]. Expressing the N protein by an AdC vector within a heterologous viral protein that is expressed on the surface of an infected cells may promote B cell responses to the N protein’s linear epitopes [29] but will likely distort the proteins’ structure and destroy conformation-dependent epitopes. Nevertheless, we obtained a strong N protein specific antibody response after the boost in hamsters that received the AdC-gDN vaccines.

The N protein as expressed by the AdC vectors induces potent and broad CD8^+^ T cell responses in mice [38]. Responses are readily detectable but modest in hamsters, which may in part relate to the paucity of hamster-specific antibodies to T cell determinants and key cytokines, which only allowed us to test for circulating CD8^+^ lymphocytes producing IFN-γ in response to the N peptide pools.

Hamsters were challenged after vaccination to assess if the vaccines protected against disease or infection. It is noteworthy that mirroring humans male hamsters were more susceptible to severe disease than female hamsters – one male hamster in the control group died and all male hamsters regardless of vaccination exhibited clinical symptoms throughout the 14-day observation period. Vaccinated female hamsters showed accelerated recovery from symptoms especially after immunization with the AdC-S-AdC-gDN combination vaccine. Both the combination vaccine and the AdC-S vaccine provided solid protection against weight loss. Despite reducing symptoms none of the vaccine regimens prevented infection; in all animals viral RNA and sgRNA were recovered at high titers from their oral cavities with only minor reductions in groups 2 and 3 on day 2 after challenge. In the same token only one group 2 animal was protected against a pulmonary infection while all other animals had detectable titers of viral RNA and sgRNA on day 4 after challenge which declined by day 14. Lung virus RNA titers on day 4 were significantly lower in groups 2 and 3 and by day 14, 3 out of 4 group 3 and 1 out of 4 group 2 animals scored negative for lung sgRNA titers while control animals and animals vaccinated only with the AdC-gDN vaccine retained titers of approximately 10^5^ copies per ml. Not only did vaccinated animals become infected but they also transmitted the virus to unvaccinated hamsters that were placed into their cages as of day 4 after challenge (data not shown). Lung pathology mirrored lung sgRNA titers and again females had less lung disease compared to males and group 3 animals tended to recover faster than group 2 animals while group 1 animals showed no difference compared to control animals although addition of the AdC-gDN vaccine reduced disease and accelerated recovery in animals that were also vaccinated with the AdC-S vaccine. By itself the AdC-gDN vaccine only induced marginal protection, which contrasts data from another group; they reported T cell-mediated protection against COVID-19 disease and a reduction in viral loads upon intravenous immunization of hamsters with a human serotype 5 Ad vector expressing the N protein [48]. Differences in results may reflect the distinct immunization protocols or differences in the challenge.

In our study the contribution of the AdC-gDN vaccine to protection induced by the AdC-S vaccines was not solely mediated by T cells, which have the advantage that upon antigen-driven activation they persist for very long periods of time [49], which would likely extend duration of protection well beyond the 4-6 months that current vaccines offer. Antibodies to the N protein strong positive correlations with relative weight of the animals and inverse correlations with disease parameters such as viral loads and lung pathology in the late euthanasia group indicating that the N protein-specific antibodies contribute to accelerated recovery from the infection.

Although available vaccines still avert serious COVID-19 disease and death, considering that we are entering the 3^rd^ year of the pandemic and the near certainty that additional variants with even more mutations that escape protection induced by current vaccines will evolve, new vaccines expressing additional SARS-CoV-2 antigens, such as the N protein to augment VNA mediated protection through additional immune mechanisms should be explored.

## MATERIAL AND METHODS

### Cell lines

HEK 293 cells were grown in Dulbecco’s Modified Eagles medium (DMEM) supplemented with 10% fetal bovine serum (FBS) and antibiotics. BHK-21/WI-2 cells were grown in DMEM supplemented with 5% FBS and antibiotics. A549-hACE-TMPRSS2 (A549A/T) cells (InvivoGen, cat no. a549-hace2tpsa) were maintained in DMEM with 10%FBS, 300µg/ml of hygromycin, 0.5 µg/ml of puromycin and antibiotics. All cells were maintained at 37°C in in a 5% CO_2_ incubator.

### Challenge virus

Challenge was conducted with the WT WAS stock, which was generated from seed stock (2019-nCoV/USA-WA1/2020), obtained from Biodefense and Emerging Infections Research (BEI) resources (Cat # NR-52281, Lot # 70036318) and expanded in Calu-3 cells at 37°C for 3 days. The stock (Lot # 12152020-1235) titers in Vero TMPRSS2 cells are 2.34×10^9^ Median Tissue Culture Infectious Dose (TCID_50_)/mL and 3.68 × 10^7^ plaque forming units (PFU)/mL, and 6 × 10^5^ PFU/mL in Vero76 cells.

### Generation and quality control of AdC vectors

Construction and expansion of the AdC-S and AdC-gDN vectors has been described [50]. Briefly the AdC-S vectors carry the gene encoding the full-length S gene (GenBank: QIC53204.1) while the AdC-gDN vectors carry the N gene (GenBank: QIC50514) that upon removal of the start and stop codons had been cloned into herpes simplex virus gD. Quality control assays for the AdC vector batches used for the hamster studies included titrations for vp content and infectious units, testing for sterility and endotoxin content and restriction enzyme digest of viral DNA, the latter to establish genetic integrity. The vectors passed the quality controls.

### Generation of VSV-S vectors

VSV vectors pseudotyped with S of SARS-CoV-2 were generated in BHK-21/WI-2 cells using a the ΔG-GFP (G*ΔG-GFP) rVSV kit (Kerafast, Boston, MA, USA) and S sequences cloned into an expression plasmid under the control of the CMV promoter as described previously [51]. One carries the same S protein as the vaccine vectors (VSV-S_SWE_). In a second the S_SWE_ sequence was modified to incorporate the E484K and N501Y RBD mutations of the B1.351 South African variant (VSV-S_SWE/B1.351_). Others express either the S protein with N501Y and P681H RBD mutations present in the B1.1.7 UK variant (VSV-S_SWE/B1.1.7_), the L452R and D614G mutations of the Indian B.1.617.2 delta variant (VSV-S _SWE/B.1.617.2_), or a full-length synthetic sequence of the omicron variant BA.1/B.1.1.529.1 (VSV-S _B.1.1.529.1_).

### Hamster challenge

Prior to challenge, hamsters were anesthetized with 80 mg/kg Ketamine and 5 mg/kg xylazine given i.m. The animals were challenged with 6 × 10^3^ PFU (Vero76 cells) in 100 μL per animal (50 μl/nostril). Administration of virus was conducted as follows: Using a calibrated P200 pipettor, 50 μl of the viral inoculum was administered dropwise into each nostril. Anesthetized animals were held upright such that the nostrils of the hamsters were pointing towards the ceiling. The tip of the pipette was placed into the first nostril and virus inoculum was slowly injected into the nasal passage, and then removed. This was repeated for the second nostril. The animal’s head was tilted back for about 20 seconds and then the animal was returned to its housing unit. ∼20 min post challenged antisedan (0.04 ml, 1mg/kg) was given i.m. to all hamsters.

### Antibody Assays

#### ELISA for anti-S1 and anti-S2 antibodies

Sera from individual hamsters were tested for S-specific antibodies by ELISA on plates coated overnight with 100 µl of a mixture of S1 and S2 (Native Antigen Company, Kidlington, UK) each diluted to 1 µg/ml in bicarbonate buffer. The next day plates were washed and blocked for 24 hr at 4°C with 150 µl of a 3% BSA-PBS solution. Sera were diluted in 3% BSA-PBS and 80 µl of the dilutions were added to the wells after the blocking solution had been discarded. Plates were incubated at room temperature for 1 hr and then washed 4 × with 150 µl of PBS. An anti-hamster IgG (H+L)-Alkaline Phosphatase antibody produced in goat (Sigma-Aldrich, St Louis, MO) diluted to 1:1000 in 3% BAS-PBS was added at 60 µl/well for 1 hour at room temperature. Plates were washed 4 × and 100 µl of substrate composed of phosphatase substrate tables (Sigma-Aldrich, St Louis, MO) diluted in 5 ml of diethanolamine buffer per tablet was added. Plates were read in an ELISA reader at 405 nm. Coated wells only receiving the substrate served to determine background. Background data were subtracted from the experimental data. Data are expressed as area under the curve for the different dilutions.

#### ELISA for RBD-binding antibodies

Sera were tested for inhibition of ACE binding to the RBD of the S protein by the Anti-SARS-CoV-2 Neutralizing Antibody Titer Serologic Assay Kit from Acro Biosystems (Newark, DE) following the manufacturer’s instructions. A positive standard with a known concentration provided by the kit was used to extrapolated anti-S antibody concentrations into µg per ml of serum.

#### Neutralization assay with S protein pseudotyped VSV vectors

VSV-S vectors was initially titrated on A549A/T cells. Cells were diluted to 4 ×10^5^ cells/ml in DMEM with 10% FBS and 5 µl of the cell dilution were added to each wells of Terasaki plates. The next day serial dilutions of VSV-S vectors were added to duplicate wells. Numbers of fluorescent cells/well were counted with a fluorescent microscope 48 hours later.

For the neutralization assay a dose of VSV-S vector that infected ∼20-40 cells/well was selected. A549A/T cells were plated into Terasaki plate wells as described above. The following day, sera were serially diluted in DMEM with 10% FBS. The VSV-S vector was diluted in medium containing 1 µg/ml of a mouse monoclonal antibody against VSV glycoprotein (sc-365019, Santa Crusz Biotechnology, Dallas, TX) and then incubated with the serum dilutions for 90 min at room temperature under gentle agitation starting with a serum dilution of 1 in 40. 5 µl aliquots of the mixtures were transferred onto Terasaki plate wells with A549A/T cells. Each serum dilution was tested in duplicates, an additional 4-6 wells were treated with VSV-S that had been incubated with medium rather than serum. Plates were incubated for 48 hours and then numbers of fluorescent cells were counted. Titers were set as the last serum dilution that reduced numbers of green-fluorescent cells by at least 50%.

#### ELISA for N protein-specific antibodies

Sera were tested for antibodies to the N protein by an ELISA on plates coated with 1 µg of a purified N protein (Abbexa, abx163973) using the same procedures as for antibodies to the S protein. A monoclonal antibody to N protein (Abbexa, MCA6373, lot# 156962) served as a positive control (data not shown).

#### T cell assays

Blood diluted in EDTA was collected 2 weeks after the 2^nd^ immunization and delivered within 6 hours after collection to the laboratory. PBMCs were purified by Ficoll^®^ Paque Plus (GE Healthcare, Chicago, IL) gradient centrifugation for 30 min at 2800 rpm. Cells were washed and seeded into 96-roundbottom well plates (0.2-1×10^6^ cells per well). Lymphocytes were stimulated with three pools of peptides representing the N sequence present in the vaccines. Peptides were 15 amino acids in length and overlapped by 10 amino acids with the adjacent peptides. Individual peptides were diluted according to the manufacturer’s instructions. For stimulation ∼10^6^ lymphocytes plated in medium containing 2% fetal calf serum and 1.5 μl/ml Golgiplug (BD Bioscience; San Jose, CA) were cultured with peptides each present at a final concentration of 2 µg/ml for 5 hr at 37°C in a 5% CO_2_ incubator. Control cells were cultured without peptides. Following stimulation cells were incubated with anti-CD8b-PE (clone eBio341, eBioscience 53-6.7 and violet live/dead dye (Thermo Fisher Scientific) at 4°C for 30 min in the dark. Cells were washed once with PBS and then fixed and permeabilized with Cytofix/Cytoperm (BD Biosciences, San Jose, CA) for 20 min. Cells were then incubated with an anti-INF-γ-FITC antibody (clone XMG1.2 BioLegend, San Diego, CA) at 4°C for 30 min in the dark. Cells were washed and fixed in 1:3 dilution of BD Cytofix fixation buffer (BD Pharmingen, San Diego CA). They were analyzed by a BD FACS Celesta (BD Biosciences, San Jose, CA) and DiVa software. Post-acquisition analyses were performed with FlowJo (TreeStar, Ashland, OR). Data shown in graph represents % of INF-γ production by CD8^+^ cells upon peptide stimulation. Background values obtained for the same cells cultured without peptide(s) were subtracted.

#### Genomic mRNA PCR Assay

The qRT-PCR assay utilizes primers and a probe specifically designed to bind to a conserved region of N gene of SARS-CoV-2. The signal is compared to a known standard curve and calculated to give copies per mL. For the qRT-PCR assay, viral RNA was first isolated from oral swab using the Qiagen MinElute virus spin kit (cat. no. 57704). For tissues it was extracted with RNA-STAT 60 (Tel-test”B”)/ chloroform, precipitated and resuspended in RNAse-free water. To generate a control for the amplification reaction, RNA was isolated from the applicable SARS-CoV-2 stock using the same procedure. The amount of RNA was determined from an O.D. reading at 260, using the estimate that 1.0 OD at A260 equals 40 µg/mL of RNA. With the number of bases known and the average base of RNA weighing 340.5 g/mole, the number of copies was then calculated, and the control diluted accordingly. A final dilution of 10^8^ copies per 3 µl was then divided into single use aliquots of 10 µl. For the master mix preparation, 2.5 mL of 2X buffer containing Taq-polymerase, obtained from the TaqMan RT-PCR kit (Bioline cat# BIO-78005), was added to a 15 mL tube. From the kit, 50 µl of the RT and 100 µl of RNAse inhibitor was added. The primer pair at 2 µM concentration was then added in a volume of 1.5 ml. Lastly, 0.5 ml of water and 350 µl of the probe at a concentration of 2 µM were added and the tube vortexed. For the reactions, 45 µl of the master mix and 5 µl of the sample RNA was added to the wells of a 96-well plate. The plates were sealed with a plastic sheet. All samples were tested in triplicate.

For control curve preparation, samples of the control RNA were prepared to contain 10^6^ to 10^7^ copies per 3 µL. Eight (8) 10-fold serial dilutions of control RNA were prepared using RNAse-free water by adding 5 µl of the control to 45 µl of water and repeating this for 7 dilutions resulting in a standard curve with a range of 1 to 10^7^ copies/reaction. Duplicate samples of each dilution were prepared as described above. If the copy number exceeded the upper detection limit, the sample was diluted as needed. For amplification, the plate was placed in an Applied Biosystems 7500 Sequence detector and amplified using the following program: 48ºC for 30 minutes, 95ºC for 10 min followed by 40 cycles of 95ºC for 15 sec, and 1 min at 55ºC. The number of copies of RNA per ml was calculated by extrapolation from the standard curve and multiplied by the reciprocal of 0.2 ml extraction volume. This gave a practical range of 50 to 5 × 10^8^ RNA copies per gram tissue.

Primers/probe sequences:

2019-nCoV_N1-F :5’-GAC CCC AAA ATC AGC GAA AT-3’

2019-nCoV_N1-R: 5’-TCT GGT TAC TGC CAG TTG AAT CTG-3’

2019-nCoV_N1-P: 5’-FAM-ACC CCG CAT TAC GTT TGG TGG ACC-BHQ1-3’.

#### Sub-genomic mRNA Assay

For the qRT-PCR assay, viral RNA is first isolated using the Qiagen MinElute virus spin kit (cat. no. 57704). For tissues, the tissue was homogenized and RNA is extracted with RNA-STAT 60 (Tel-test”B”)/chloroform, precipitated and resuspended in AVE Buffer (Qiagen 1020953). The qRT-PCR assay utilizes primers and a probe specifically designed to amplify and bind to a conserved region of N gene of SARS-CoV-2. The signal was compared to a known standard curve and calculated to give copies per ml. To generate a control for the amplification reaction, RNA was isolated from the applicable virus stock using the same procedure. The amount of viral RNA was determined comparing it to a known quantity of plasmid control. A final dilution of 10^8^ copies per 3 µl was then divided into single use aliquots of 10 µl. For the master mix preparation, 2.5 ml of 2X buffer containing Taq-polymerase, obtained from the TaqMan RT-PCR kit (Bioline #BIO-78005), was added to a 15 ml tube. From the kit, 50 µl of the RT and 100 µl of RNAse inhibitor was also added. The primer pair at 2 µM concentration was then added in a volume of 1.5 ml. Lastly, 0.5 ml of water and 350 µl of the probe at a concentration of 2 µM are added and the tube vortexed. For the reactions, 45 µl of the master mix and 5 µl of the sample RNA were added to the wells of a 96-well plate. The plates were sealed with a plastic sheet. All samples were tested in triplicate. For control curve preparation, the control viral RNA was prepared to contain 10^6^ to 10^7^ copies per 3 µl. Seven 10-fold serial dilutions of control RNA were prepared using RNAse-free water by adding 5 µl of the control to 45 µl of water and repeating this for 7 dilutions. This gives a standard curve with a range of 1 to 10^7^ copies/reaction. The sub-genomic-N uses a known plasmid for its curve. Duplicate samples of each dilution were prepared as described above. If the copy number exceeded the upper detection limit, the sample was diluted as needed. For amplification, the plate was placed in an Applied Biosystems 7500 Sequence detector and amplified using the following program: 48°C for 30 minutes, 95°C for 10 min followed by 40 cycles of 95 °C for 15 sec, and 1 min at 55°C. The number of copies of RNA per ml is calculated by extrapolation from the standard curve and multiplying by the reciprocal of 0.2 ml extraction volume. This gives a practical range of 50 to 5 × 10^8^ RNA copies per mL for oral swabs and BAL, and for lung tissue the viral loads are given per gram.

sg-N-F: 5’-CGATCTCTTGTAGATCTGTTCTC-3’

sg-N-R: 5’-GGTGAACCAAGACGCAGTAT-3’

Sg-N-P: 5’-6-FAM/TAACCAGAA/ZEN/TGGAGAACGCAGTGGG/3IABkFQ/

#### Lung Histology

Parts of the lungs were placed in 10% neutral buffered formalin for histopathologic analysis. Lung was processed to hematoxylin and eosin (H&E) stained slides and examined by a board-certified pathologist at Experimental Pathology Laboratories, Inc. (EPL^®^) in Sterling, Virginia. Findings were graded from one to five, depending upon severity.

#### Statistical analyses

Data were analyzed by 2-or 1-way ANOVA or Kruskal-Wallis test with Dunn correction for multiple comparisons. Correlations were carried out by one-tailed Spearmanʼs rank correlation tests. *P* values equal or less than 0.05 were considered significant. The analyses were carried out by GraphPad Prism.

## Acknowledgements

This work was funded by grants from The G. Harold and Leila Y. Mathers Charitable Foundation, the Commonwealth of Pennsylvania, and the Wistar Science Discovery Fund. MH was the recipient of fellowship from Fellowship from Janssen Scientific Affairs. Support for Shared Resources utilized in this study was provided by Cancer Center Support Grant (CCSG) P30CA010815 to The Wistar Institute. We wish to thank Dr. Elledge (Massachusetts General Hospital, Boston, MA) for providing the cDNA sequences for N and S of SARS-CoV-2.

---

**Supplemental Figure 1. Weight loss and lung pathology**

The two graphs show relative weights over time after challenge in female and male hamsters. [B] The graphs show representative lung sections from animals of groups 1-4 collected on day 4 or 14 after challenge. Some of the diseased areas are highlighted by arrows.

**Supplemental Figure 2. Mutations within S protein that affect HLA class I epitopes**

Protein sequence of S is shown on top of each line, -: conserved sequences, letters in red: mutations, − deletions, empty spaces: insertion within one of the sequences, MHC class I epitopes highlighted; grey: score ≥0.9, underlined: score 0.8 -<0.9.

**Supplemental Figure 3. Mutations within S protein that affect HLA class II epitopes**

Protein sequence of Sis shown on top of each line, -: conserved sequences, letters in red: mutations, − deletions, empty spaces: insertion within one of the other sequences, MHC class II epitopes highlighted: grey: adjusted rank <0.5, underlined: adjusted range ≥0.5-1.

**Supplemental Figure 4. Mutations within N protein that affect HLA class I epitopes**

Protein sequence of N is shown on top of each line, -: conserved sequences, letters in red: mutations, − deletions, empty spaces: insertion within one of the sequences, MHC class I epitopes highlighted; grey: score ≥0.9, underlined: score 0.8 -<0.9.

**Supplemental Figure 5. Mutations within N protein that affect HLA class II epitopes**

Protein sequence of N is shown on top of each line., -: conserved sequences, letters in red: mutations, − – deletions, empty spaces: insertion within one of the other sequences, MHC class II epitopes highlighted: grey: adjusted rank <0.5, underlined: adjusted range ≥0.5-1.

**Supplemental Figure 6. Mutations within S protein that affect linear B cell epitopes**

Protein sequence of S is shown on top of each line, -: conserved sequences, letters in red: mutations, − deletions, empty spaces: insertion within one of the sequences, published linear B cell epitopes are highlighted.

**Supplemental Figure 7. Mutations within N protein that affect linear B cell epitopes**

Protein sequence of N is shown on top of each line, -: conserved sequences, letters in red: mutations, − deletions, empty spaces: insertion within one of the sequences, published linear B cell epitopes are highlighted.

