## Supplemental figures for "Prime-boost vaccinations with two serologically distinct chimpanzee adenovirus vectors expressing SARS-CoV-2 spike or nucleocapsid tested in a hamster COVID-19 model"

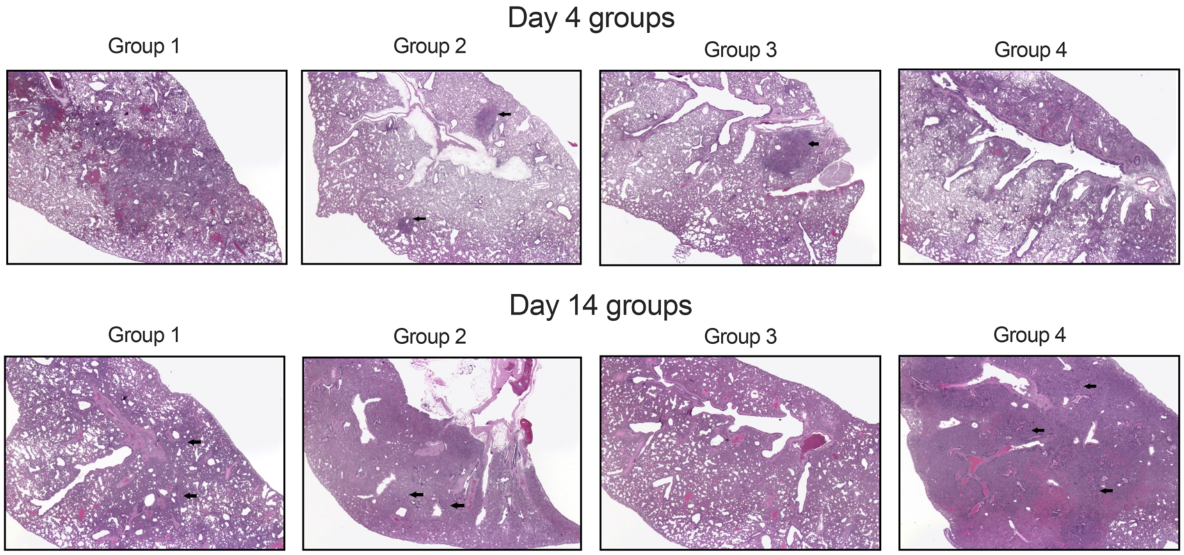


**Supplemental Figure 1. Weight loss and lung pathology**
[A] The two graphs show relative weights over time after challenge in female and male hamsters. [B] The graphs show representative lung sections from animals of groups 1-4 collected on day 4 or 14 after challenge.  Some of the diseased areas are highlighted by arrows.

SARS-CoV-2 spike protein sequences and HLA class I epitopes

```
Wild-type     1  MFVFLVLLPLVSSQCVNLTTRTQLPPAYTNSFTRGVYYPDKVFRSSVLHSTQDLFLPFFSNVTWFHAIHVSGTNGTK
Alpha            --------------------------------------------------------------------ΔΔ-------
Beta             -----------------F-----------------------------------------------------------
Delta            ------------------R----------------------------------------------------------
Omicron          ------------------------------------------------------------------V-ΔΔ-------

Wild-type    78  RFDNPVLPFNDGVYFASTEKSNIIRGWIFGTTLDSKTQSLLIVNNATNVVIKVCEFQFCNDPFLGVYYHKNNKSWME
Alpha            ------------------------------------------------------------------Δ----------
Beta             --A--------------------------------------------------------------------------
Delta            ----------------------------------------------------------------D------------
Omicron          -----------------I----------------------------------------------DΔΔΔ---------

Wild-type   155  SEFRVYSSANNCTFEYVSQPFLMDLEGKQGNFKNLREFVFKNIDGYFKIYSKHTPINLVR   DLPQGFSALEPLVD
Alpha            ------------------------------------------------------------   --------------
Beta             ------------------------------------------------------------   G-------------
Delta            -ΔΔG--------------------------------------------------------   --------------
Omicron          --------------------------------------------------------ΔI--EPE--------------

Wild-type   229   LPIGINITRFQTLLALHRSYLTPGDSSSGWTAGAAAYYVGYLQPRTFLLKYNENGTITDAVDCALDPLSETKCTLKS
Alpha            -----------------------------------------------------------------------------
Beta             -------------ΔΔΔ-------------------------------------------------------------
Delta            -----------------------------------------------------------------------------
Omicron          -----------------------------------------------------------------------------

Wild-type   306  FTVEKGIYQTSNFRVQPTESIVRFPNITNLCPFGEVFNATRFASVYAWNRKRISNCVADYSVLYNSASFSTFKCYGV
Alpha            -----------------------------------------------------------------------------
Beta             -----------------------------------------------------------------------------
Delta            -----------------------------------------------------------------------------
Omicron          ---------------------------------D-------------------------------L-P-F-------

Wild-type   382  SPTKLNDLCFTNVYADSFVIRGDEVRQIAPGQTGKIADYNYKLPDDFTGCVIAWNSNNLDSKVGGNYNYLYRLFRKS
Alpha            -----------------------------------------------------------------------------
Beta             ----------------------------------N------------------------------------------
Delta            ---------------------------------------------------------------------R-------
Omicron          ----------------------------------N----------------------K-----S-------------

Wild-type   460  NLKPFERDISTEIYQAGSTPCNGVEGFNCYFPLQSYGFQPTNGVGYQPYRVVVLSFELLHAPATVCGPKKSTNLVKN
Alpha            -----------------------------------------------------------------------------
Beta             ------------------------K----------------Y-----------------------------------
Delta            ------------------K----------------------------------------------------------
Omicron          -----------------NK-----A--------R--S-R--Y---H-------------------------------

Wild-type   537  KCVNFNFNGLTGTGVLTESNKKFLPFQQFGRDIADTTDAVRDPQTLEILDITPCSFGGVSVITPGTNTSNQVAVLYQ
Alpha            ---------------------------------D-------------------------------------------
Beta             -----------------------------------------------------------------------------
Delta            -----------------------------------------------------------------------------
Omicron          ----------K------------------------------------------------------------------

Wild-type   614  DVNCTEVPVAIHADQLTPTWRVYSTGSNVFQTRAGCLIGAEHVNNSYECDIPIGAGICASYQTQTNSPRRARSVASQ
Alpha            G------------------------------------------------------------------H---------
Beta             -----------------------------------------------------------------------------
Delta            G------------------------------------------------------------------R---------
Omicron          G----------------------------------------Y-----------------------K-H---------

Wild-type   691  SIIAYTMSLGAENSVAYSNNSIAIPTNFTISVTTEILPVSMTKTSVDCTMYICGDSTECSNLLLQYGSFCTQLNRAL
Alpha            -------------------------I---------------------------------------------------
Beta             ----------V------------------------------------------------------------------
Delta            -----------------------------------------------------------------------------
Omicron          -------------------------------------------------------------------------K---

Wild-type   768  TGIAVEQDKNTQEVFAQVKQIYKTPPIKDCGGFNFSQILPDPSKPSKRSFIEDLLFNKVTLADAGFIKQYGDCLGDI
Alpha            -----------------------------------------------------------------------------
Beta             -----------------------------------------------------------------------------
Delta            -----------------------------------------------------------------------------
Omicron          ----------------------------Y------------------------------------------------

Wild-type   845  AARDLICAQKFNGLTVLPPLLTDEMIAQYTSALLAGTITSGWTFGAGAALQIPFAMQMAYRFNGIGVTQNVLYENQK
Alpha            -----------------------------------------------------------------------------
Beta             -----------------------------------------------------------------------------
Delta            -----------------------------------------------------------------------------
Omicron          -----------K-----------------------------------------------------------------

Wild-type   922  LIANQFNSAIGKIQDSLSSTASALGKLQDVVNQNAQALNTLVKQLSSNFGAISSVLNDILSRLDKVEAEVQIDRLIT
Alpha            ----------------------------G------------------------------------------------
Beta             ----------------------------G------------------------------------------------
Delta            -----------------------------------------------------------------------------
Omicron          --------------------------------H--------------K-----------F-----------------

Wild-type   999  GRLQSLQTYVTQQLIRAAEIRASANLAATKMSECVLGQSKRVDFCGKGYHLMSFPQSAPHGVVFLHVTYVPAQEKNF
Alpha            -----------------------------------------------------------------------------
Beta             -----------------------------------------------------------------------------
Delta            -----------------------------------------------------------------------------
Omicron          -----------------------------------------------------------------------------

Wild-type  1076  TTAPAICHDGKAHFPREGVFVSNGTHWFVTQRNFYEPQIITTDNTFVSGNCDVVIGIVNNTVYDPLQPELDSFKEEL
Alpha            -----------------------------------------------------------------------------
Beta             -----------------------------------------------------------------------------
Delta            -----------------------------------------------------------------------------
Omicron          -----------------------------------------------------------------------------

Wild-type  1153  DKYFKNHTSPDVDLGDISGINASVVNIQKEIDRLNEVAKNLNESLIDLQELGKYEQYIKWPWYIWLGFIAGLIAIVM
Alpha            -----------------------------------------------------------------------------
Beta             -----------------------------------------------------------------------------
Delta            -----------------------------------------------------------------------------
Omicron          -----------------------------------------------------------------------------

Wild-type  1230  VTIMLCCMTSCCSCLKGCCSCGSCCKFDEDDSEPVLKGVKLHYT
Alpha            --------------------------------------------
Beta             --------------------------------------------
Delta            --------------------------------------------
Omicron          --------------------------------------------
```

**Supplemental Figure 2. Mutations within S protein that affect HLA class I epitopes**

Protein sequence of S is shown on top of each line, - : conserved sequences, letters in red: mutations, D – deletions, empty spaces: insertion within one of the sequences, MHC class I epitopes highlighted; grey: score ≥0.9, underlined: score 0.8 - <0.9.

SARS-CoV-2 spike protein sequences and HLA class II epitopes

```
Wild-type     1  MFVFLVLLPLVSSQCVNLTTRTQLPPAYTNSFTRGVYYPDKVFRSSVLHSTQDLFLPFFSNVTWFHAIHVSGTNGTK
Alpha            --------------------------------------------------------------------ΔΔ-------
Beta             -----------------F-----------------------------------------------------------
Delta            ------------------R----------------------------------------------------------
Omicron          ------------------------------------------------------------------V-ΔΔ-------

Wild-type    78  RFDNPVLPFNDGVYFASTEKSNIIRGWIFGTTLDSKTQSLLIVNNATNVVIKVCEFQFCNDPFLGVYYHKNNKSWME
Alpha            ------------------------------------------------------------------Δ----------
Beta             --A--------------------------------------------------------------------------
Delta            ----------------------------------------------------------------D------------
Omicron          -----------------I----------------------------------------------DΔΔΔ---------

Wild-type   155  SEFRVYSSANNCTFEYVSQPFLMDLEGKQGNFKNLREFVFKNIDGYFKIYSKHTPINLVR   DLPQGFSALEPLVD
Alpha            ------------------------------------------------------------   --------------
Beta             ------------------------------------------------------------   G-------------
Delta            -ΔΔG--------------------------------------------------------   --------------
Omicron          --------------------------------------------------------ΔI--EPE--------------

Wild-type   229  LPIGINITRFQTLLALHRSYLTPGDSSSGWTAGAAAYYVGYLQPRTFLLKYNENGTITDAVDCALDPLSETKCTLKS
Alpha            -----------------------------------------------------------------------------
Beta             -------------ΔΔΔ-------------------------------------------------------------
Delta            -----------------------------------------------------------------------------
Omicron          -----------------------------------------------------------------------------

Wild-type   306  FTVEKGIYQTSNFRVQPTESIVRFPNITNLCPFGEVFNATRFASVYAWNRKRISNCVADYSVLYNSASFSTFKCYGV
Alpha            -----------------------------------------------------------------------------
Beta             -----------------------------------------------------------------------------
Delta            -----------------------------------------------------------------------------
Omicron          ---------------------------------D-------------------------------L-P-F-------

Wild-type   382  SPTKLNDLCFTNVYADSFVIRGDEVRQIAPGQTGKIADYNYKLPDDFTGCVIAWNSNNLDSKVGGNYNYLYRLFRKS
Alpha            -----------------------------------------------------------------------------
Beta             ----------------------------------N------------------------------------------
Delta            ---------------------------------------------------------------------R-------
Omicron          ----------------------------------N----------------------K-----S-------------

Wild-type   460  NLKPFERDISTEIYQAGSTPCNGVEGFNCYFPLQSYGFQPTNGVGYQPYRVVVLSFELLHAPATVCGPKKSTNLVKN
Alpha            -----------------------------------------Y-----------------------------------
Beta             ------------------------K----------------Y-----------------------------------
Delta            ------------------K----------------------------------------------------------
Omicron          -----------------NK-----A--------R--S-R--Y-----------------------------------

Wild-type   537  KCVNFNFNGLTGTGVLTESNKKFLPFQQFGRDIADTTDAVRDPQTLEILDITPCSFGGVSVITPGTNTSNQVAVLYQ
Alpha            ---------------------------------D-------------------------------------------
Beta             -----------------------------------------------------------------------------
Delta            -----------------------------------------------------------------------------
Omicron          -----------------------------------------------------------------------------

Wild-type   614  DVNCTEVPVAIHADQLTPTWRVYSTGSNVFQTRAGCLIGAEHVNNSYECDIPIGAGICASYQTQTNSPRRARSVASQ
Alpha            G------------------------------------------------------------------H---------
Beta             -----------------------------------------------------------------------------
Delta            G------------------------------------------------------------------R---------
Omicron          G-----------------------------------------Y-----------------------K-H---------

Wild-type   691  SIIAYTMSLGAENSVAYSNNSIAIPTNFTISVTTEILPVSMTKTSVDCTMYICGDSTECSNLLLQYGSFCTQLNRAL
Alpha            -------------------------I---------------------------------------------------
Beta             ----------V------------------------------------------------------------------
Delta            -----------------------------------------------------------------------------
Omicron          -------------------------------------------------------------------------K---

Wild-type   768  TGIAVEQDKNTQEVFAQVKQIYKTPPIKDCGGFNFSQILPDPSKPSKRSFIEDLLFNKVTLADAGFIKQYGDCLGDI
Alpha            -----------------------------------------------------------------------------
Beta             -----------------------------------------------------------------------------
Delta            -----------------------------------------------------------------------------
Omicron          ----------------------------Y------------------------------------------------

Wild-type   845  AARDLICAQKFNGLTVLPPLLTDEMIAQYTSALLAGTITSGWTFGAGAALQIPFAMQMAYRFNGIGVTQNVLYENQK
Alpha            -----------------------------------------------------------------------------
Beta             -----------------------------------------------------------------------------
Delta            -----------------------------------------------------------------------------
Omicron          -----------K-----------------------------------------------------------------

Wild-type   922  LIANQFNSAIGKIQDSLSSTASALGKLQDVVNQNAQALNTLVKQLSSNFGAISSVLNDILSRLDKVEAEVQIDRLIT
Alpha            ----------------------------G------------------------------------------------
Beta             ----------------------------G------------------------------------------------
Delta            -----------------------------------------------------------------------------
Omicron          --------------------------------H--------------K-----------F-----------------

Wild-type   999  GRLQSLQTYVTQQLIRAAEIRASANLAATKMSECVLGQSKRVDFCGKGYHLMSFPQSAPHGVVFLHVTYVPAQEKNF
Alpha            -----------------------------------------------------------------------------
Beta             -----------------------------------------------------------------------------
Delta            -----------------------------------------------------------------------------
Omicron          -----------------------------------------------------------------------------

Wild-type  1076  TTAPAICHDGKAHFPREGVFVSNGTHWFVTQRNFYEPQIITTDNTFVSGNCDVVIGIVNNTVYDPLQPELDSFKEEL
Alpha            -----------------------------------------------------------------------------
Beta             -----------------------------------------------------------------------------
Delta            -----------------------------------------------------------------------------
Omicron          -----------------------------------------------------------------------------
Wild-type  1153  DKYFKNHTSPDVDLGDISGINASVVNIQKEIDRLNEVAKNLNESLIDLQELGKYEQYIKWPWYIWLGFIAGLIAIVM
Alpha            -----------------------------------------------------------------------------
Beta             -----------------------------------------------------------------------------
Delta            -----------------------------------------------------------------------------
Omicron          -----------------------------------------------------------------------------
Wild-type  1230  VTIMLCCMTSCCSCLKGCCSCGSCCKFDEDDSEPVLKGVKLHYT
Alpha            --------------------------------------------
Beta             --------------------------------------------
Delta            --------------------------------------------
Omicron          --------------------------------------------
```

**Supplemental Figure 3. Mutations within S protein that affect HLA class II epitopes**

Protein sequence of Sis shown on top of each line, - : conserved sequences, letters in red: mutations, D – deletions, empty spaces: insertion within one of the other sequences, MHC class II epitopes highlighted: grey: adjusted rank <0.5, underlined: adjusted range ≥0.5-1.

SARS-CoV-2 nucleoapsid protein sequences and HLA class I epitopes

```
Wild-type   1   MSDNGPQNQRNAPRITFGGPSDSTGSNQNGERSGARSKQRRPQGLPNNTASWFTALTQHGKEDLKFPRG
Alpha           --L------------------------------------------------------------------
Beta            ---------------------------------------------------------------------
Delta           ---------------------------------------------------------------------
Omicron         ------------L--------------------------------------------------------

Wild-type   70  QGVPINTNSSPDDQIGYYRRATRRIRGGDGKMKDLSPRWYFYYLGTGPEAGLPYGANKDGIIWVATEGA
Alpha           ---------------------------------------------------------------------
Beta            ---------------------------------------------------------------------
Delta           ---------------------------------------------------------------------
Omicron         ---------------------------------------------------------------------

Wild-type   139 LNTPKDHIGTRNPANNAAIVLQLPQGTTLPKGFYAEGSRGGSQASSRSSSRSRNSSRNSTPGSNRGTSP
Alpha           ---------------------------------------------------------------------
Beta            ------------------------------------------------------------------I--
Delta           ----------------------------------------------------------------M----
Omicron         ----------------------------------------------------------------KR---

Wild-type   208 ARMAGNGGDAALALLLLDRLNQLESKMSGKGQQQQGQTVTKKSAAEASKKPRQKRTATKAYNVTQAFGR
Alpha           ---------------------------F-----------------------------------------
Beta            ---------------------------------------------------------------------
Delta           ---------------------------------------------------------------------
Omicron         ---------------------------------------------------------------------

Wild-type   277 RGPEQTQGNFGDQELIRQGTDYKHWPQIAQFAPSASAFFGMSRIGMEVTPSGTWLTYTGAIKLDDKDPN
Alpha           ---------------------------------------------------------------------
Beta            ---------------------------------------------------------------------
Delta           ---------------------------------------------------------------------
Omicron         ---------------------------------------------------------------------

Wild-type   346 FKDQVILLNKHIDAYKTFPPTEPKKDKKKKADETQALPQRQKKQQTVTLLPAADLDDFSKQLQQSMSSA
Alpha           ---------------------------------------------------------------------
Beta            ---------------------------------------------------------------------
Delta           -------------------------------Y-------------------------------------
Omicron         ---------------------------------------------------------------------

Wild-type   415 DSTQA
Alpha           -----
Beta            -----
Delta           -----
Omicron         -----
```

**Supplemental Figure 4. Mutations within N protein that affect HLA class I epitopes**

Protein sequence of N is shown on top of each line, - : conserved sequences, letters in red: mutations, D – deletions, empty spaces: insertion within one of the sequences, MHC class I epitopes highlighted; grey: score ≥0.9, underlined: score 0.8 - <0.9.

SARS-CoV-2 nucleocapsid protein sequences and HLA class II epitopes

```
Wild-type    1  MSDNGPQNQRNAPRITFGGPSDSTGSNQNGERSGARSKQRRPQGLPNNTASWFTALTQHGKEDLKFPRG
Alpha           --L------------------------------------------------------------------
Beta            ---------------------------------------------------------------------
Delta           ---------------------------------------------------------------------
Omicron         ------------L--------------------------------------------------------

Wild-type   70  QGVPINTNSSPDDQIGYYRRATRRIRGGDGKMKDLSPRWYFYYLGTGPEAGLPYGANKDGIIWVATEGA
Alpha           ---------------------------------------------------------------------
Beta            ---------------------------------------------------------------------
Delta           ---------------------------------------------------------------------
Omicron         ---------------------------------------------------------------------

Wild-type  139  LNTPKDHIGTRNPANNAAIVLQLPQGTTLPKGFYAEGSRGGSQASSRSSSRSRNSSRNSTPGSNRGTSP
Alpha           ---------------------------------------------------------------------
Beta            ------------------------------------------------------------------I--
Delta           ----------------------------------------------------------------M----
Omicron         ----------------------------------------------------------------KR---

Wild-type  208  ARMAGNGGDAALALLLLDRLNQLESKMSGKGQQQQGQTVTKKSAAEASKKPRQKRTATKAYNVTQAFGR
Alpha           ---------------------------F-----------------------------------------
Beta            ---------------------------------------------------------------------
Delta           ---------------------------------------------------------------------
Omicron         ---------------------------------------------------------------------

Wild-type  277  RGPEQTQGNFGDQELIRQGTDYKHWPQIAQFAPSASAFFGMSRIGMEVTPSGTWLTYTGAIKLDDKDPN
Alpha           ---------------------------------------------------------------------
Beta            ---------------------------------------------------------------------
Delta           ---------------------------------------------------------------------
Omicron         ---------------------------------------------------------------------

Wild-type  346  FKDQVILLNKHIDAYKTFPPTEPKKDKKKKADETQALPQRQKKQQTVTLLPAADLDDFSKQLQQSMSSA
Alpha           ---------------------------------------------------------------------
Beta            ---------------------------------------------------------------------
Delta           -------------------------------Y-------------------------------------
Omicron         ---------------------------------------------------------------------

Wild-type  415  DSTQA
Alpha           -----
Beta            -----
Delta           -----
Omicron         -----
```

**Supplemental Figure 5. Mutations within N protein that affect HLA class II epitopes**

Protein sequence of N is shown on top of each line. , - : conserved sequences, letters in red: mutations, D – deletions, empty spaces: insertion within one of the other sequences, MHC class II epitopes highlighted: grey: adjusted rank <0.5, underlined: adjusted range ≥0.5-1.

SARS-CoV-2 spike protein sequences and linear B cell epitopes

```
Wild-type     1  MFVFLVLLPLVSSQCVNLTTRTQLPPAYTNSFTRGVYYPDKVFRSSVLHSTQDLFLPFFSNVTWFHAIHVSGTNGTK
Alpha            --------------------------------------------------------------------ΔΔ-------
Beta             -----------------F-----------------------------------------------------------
Delta            ------------------R----------------------------------------------------------
Omicron          ------------------------------------------------------------------V-ΔΔ-------
Wild-type    78  RFDNPVLPFNDGVYFASTEKSNIIRGWIFGTTLDSKTQSLLIVNNATNVVIKVCEFQFCNDPFLGVYYHKNNKSWME
Alpha            ------------------------------------------------------------------Δ----------
Beta             --A--------------------------------------------------------------------------
Delta            ----------------------------------------------------------------D------------
Omicron          -----------------I----------------------------------------------DΔΔΔ---------
Wild-type   155  SEFRVYSSANNCTFEYVSQPFLMDLEGKQGNFKNLREFVFKNIDGYFKIYSKHTPINLVR   DLPQGFSALEPLVD
Alpha            ------------------------------------------------------------   --------------
Beta             ------------------------------------------------------------   G-------------
Delta            -ΔΔG--------------------------------------------------------   --------------
Omicron          --------------------------------------------------------ΔI--EPE--------------
Wild-type   229  LPIGINITRFQTLLALHRSYLTPGDSSSGWTAGAAAYYVGYLQPRTFLLKYNENGTITDAVDCALDPLSETKCTLKS
Alpha            -----------------------------------------------------------------------------
Beta             -------------ΔΔΔ-------------------------------------------------------------
Delta            -----------------------------------------------------------------------------
Omicron          -----------------------------------------------------------------------------
Wild-type   306  FTVEKGIYQTSNFRVQPTESIVRFPNITNLCPFGEVFNATRFASVYAWNRKRISNCVADYSVLYNSASFSTFKCYGV
Alpha            ----------------------------------------------------------------------------
Beta             ----------------------------------------------------------------------------
Delta            ----------------------------------------------------------------------------
Omicron          ---------------------------------D-------------------------------L-P-F------
Wild-type   382  SPTKLNDLCFTNVYADSFVIRGDEVRQIAPGQTGKIADYNYKLPDDFTGCVIAWNSNNLDSKVGGNYNYLYRLFRKS
Alpha            ------------------------------------------------------------------------------
Beta             -----------------------------------N------------------------------------------
Delta            ----------------------------------------------------------------------R-------
Omicron          -----------------------------------N----------------------K-----S-------------
Wild-type   460  NLKPFERDISTEIYQAGSTPCNGVEGFNCYFPLQSYGFQPTNGVGYQPYRVVVLSFELLHAPATVCGPKKSTNLVKN
Alpha            -----------------------------------------------------------------------------
Beta             ------------------------K----------------Y-----------------------------------
Delta            ------------------K----------------------------------------------------------
Omicron          -----------------NK-----A--------R--S-R--Y---H-------------------------------
Wild-type   537  KCVNFNFNGLTGTVLTESNKKFLPFQQFGRDIADTTDAVRDPQTLEILDITPCSFGGVSVITPGTNTSNQVAVLYQ
Alpha            ---------------------------------D-------------------------------------------
Beta             -----------------------------------------------------------------------------
Delta            -----------------------------------------------------------------------------
Omicron          ----------K------------------------------------------------------------------
Wild-type   614  DVNCTEVPVAIHADQLTPTWRVYSTGSNVFQTRAGCLIGAEHVNNSYECDIPIGAGICASYQTQTNSPRRARSVASQ
Alpha            G------------------------------------------------------------------H---------
Beta             -----------------------------------------------------------------------------
Delta            G------------------------------------------------------------------R---------
Omicron          G----------------------------------------Y-----------------------K-H---------
Wild-type   691  SIIAYTMSLGAENSVAYSNNSIAIPTNFTISVTTEILPVSMTKTSVDCTMYICGDSTECSNLLLQYGSFCTQLNRAL
Alpha            -------------------------I---------------------------------------------------
Beta             ----------V------------------------------------------------------------------
Delta            -----------------------------------------------------------------------------
Omicron          -------------------------------------------------------------------------K---
Wild-type   768  TGIAVEQDKNTQEVFAQVKQIYKTPPIKDCGGFNFSQILPDPSKPSKRSFIEDLLFNKVTLADAGFIKQYGDCLGDI
Alpha            -----------------------------------------------------------------------------
Beta             -----------------------------------------------------------------------------
Delta            -----------------------------------------------------------------------------
Omicron          ----------------------------Y------------------------------------------------
Wild-type   845  AARDLICAQKFNGLTVLPPLLTDEMIAQYTSALLAGTITSGWTFGAGAALQIPFAMQMAYRFNGIGVTQNVLYENQK
Alpha            -----------------------------------------------------------------------------
Beta             -----------------------------------------------------------------------------
Delta            -----------------------------------------------------------------------------
Omicron          -----------K-----------------------------------------------------------------
Wild-type   922  LIANQFNSAIGKIQDSLSSTASALGKLQDVVNQNAQALNTLVKQLSSNFGAISSVLNDILSRLDKVEAEVQIDRLIT
Alpha            ----------------------------G------------------------------------------------
Beta             ----------------------------G------------------------------------------------
Delta            -----------------------------------------------------------------------------
Omicron          --------------------------------H--------------K-----------F-----------------
Wild-type   999  GRLQSLQTYVTQQLIRAAEIRASANLAATKMSECVLGQSKRVDFCGKGYHLMSFPQSAPHGVVFLHVTYVPAQEKNF
Alpha            -----------------------------------------------------------------------------
Beta             -----------------------------------------------------------------------------
Delta            -----------------------------------------------------------------------------
Omicron          -----------------------------------------------------------------------------
Wild-type  1076  TTAPAICHDGKAHFPREGVFVSNGTHWFVTQRNFYEPQIITTDNTFVSGNCDVVIGIVNNTVYDPLQPELDSFKEEL
Alpha            -----------------------------------------------------------------------------
Beta             -----------------------------------------------------------------------------
Delta            -----------------------------------------------------------------------------
Omicron          -----------------------------------------------------------------------------
Wild-type  1153  DKYFKNHTSPDVDLGDISGINASVVNIQKEIDRLNEVAKNLNESLIDLQELGKYEQYIKWPWYIWLGFIAGLIAIVM
Alpha            -----------------------------------------------------------------------------
Beta             -----------------------------------------------------------------------------
Delta            -----------------------------------------------------------------------------
Omicron          -----------------------------------------------------------------------------
Wild-type  1230  VTIMLCCMTSCCSCLKGCCSCGSCCKFDEDDSEPVLKGVKLHYT
Alpha            --------------------------------------------
Beta             --------------------------------------------
Delta            --------------------------------------------
Omicron          --------------------------------------------
```

**Supplemental Figure 6. Mutations within S protein that affect linear B cell epitopes**

Protein sequence of S is shown on top of each line, - : conserved sequences, letters in red: mutations, D – deletions, empty spaces: insertion within one of the sequences, published linear B cell epitopes are highlighted.

SARS-CoV-2 nucleocapsid protein sequences and linear B cell epitopes

```
Wild-type     1  MSDNGPQNQRNAPRITFGGPSDSTGSNQNGERSGARSKQRRPQGLPNNTASWFTALTQHGKEDLKFPRG
Alpha            --L------------------------------------------------------------------
Beta             ---------------------------------------------------------------------
Delta            ---------------------------------------------------------------------
Omicron          ------------L--------------------------------------------------------

Wild-type    70  QGVPINTNSSPDDQIGYYRRATRRIRGGDGKMKDLSPRWYFYYLGTGPEAGLPYGANKDGIIWVATEGA
Alpha            ---------------------------------------------------------------------
Beta             ---------------------------------------------------------------------
Delta            ---------------------------------------------------------------------
Omicron          ---------------------------------------------------------------------

Wild-type   139  LNTPKDHIGTRNPANNAAIVLQLPQGTTLPKGFYAEGSRGGSQASSRSSSRSRNSSRNSTPGSNRGTSP
Alpha            ---------------------------------------------------------------------
Beta             ------------------------------------------------------------------I--
Delta            ----------------------------------------------------------------M----
Omicron          ----------------------------------------------------------------KR---

Wild-type   208  ARMAGNGGDAALALLLLDRLNQLESKMSGKGQQQQGQTVTKKSAAEASKKPRQKRTATKAYNVTQAFGR
Alpha            ---------------------------F-----------------------------------------
Beta             ---------------------------------------------------------------------
Delta            ---------------------------------------------------------------------
Omicron          ---------------------------------------------------------------------

Wild-type   277  RGPEQTQGNFGDQELIRQGTDYKHWPQIAQFAPSASAFFGMSRIGMEVTPSGTWLTYTGAIKLDDKDPN
Alpha            ---------------------------------------------------------------------
Beta             ---------------------------------------------------------------------
Delta            ---------------------------------------------------------------------
Omicron          ---------------------------------------------------------------------

Wild-type   346  FKDQVILLNKHIDAYKTFPPTEPKKDKKKKADETQALPQRQKKQQTVTLLPAADLDDFSKQLQQSMSSA
Alpha            ---------------------------------------------------------------------
Beta             ---------------------------------------------------------------------
Delta            -------------------------------Y-------------------------------------
Omicron          ---------------------------------------------------------------------

Wild-type   415  DSTQA
Alpha            -----
Beta             -----
Delta            -----
Omicron          -----
```

**Supplemental Figure 7. Mutations within N protein that affect linear B cell epitopes**
Protein sequence of N is shown on top of each line, - : conserved sequences, letters in red: mutations, D – deletions, empty spaces: insertion within one of the sequences, published linear B cell epitopes are highlighted.
